# *spe-43* is required for sperm activation in *C. elegans*

**DOI:** 10.1101/206441

**Authors:** Amber R. Krauchunas, Ernesto Mendez, Julie Zhouli Ni, Marina Druzhinina, Amanda Mulia, Jean Parry, Sam Guoping Gu, Gillian M. Stanfield, Andrew Singson

**Affiliations:** Waksman Institute of Microbiology and Department of Genetics, Rutgers University, Piscataway, NJ, USA; Department of Molecular Biology and Biochemistry, Rutgers University, Piscataway, NJ, USA; Department of Human Genetics, University of Utah, Salt Lake City, UT, USA

**Author notes:** Present address: Department of Biology, Georgian Court University, Lakewood, NJ, USA. **Corresponding author contact information:** Amber R. Krauchunas, Waksman Institute of Microbiology, 190 Frelinghuysen Rd., Rutgers University, Piscataway, NJ 08854, USA.

**Keywords:** Sperm activation, spermiogenesis, sperm, *C. elegans*

## Abstract

Successful fertilization requires that sperm are activated prior to contacting an oocyte. In *C. elegans*, this activation process, called spermiogenesis, transforms round immobile spermatids into motile, fertilization-competent spermatozoa. We describe the phenotypic and genetic characterization of *spe-43*, a new component of the *spe-8* pathway, which is required for spermiogenesis in hermaphrodites; *spe-43* hermaphrodites are self-sterile, while *spe-43* males show wild-type fertility. When exposed to Pronase to activate sperm *in vitro*, *spe-43* spermatids form long rigid spikes radiating outward from the cell periphery instead of forming a motile pseudopod, indicating that spermiogenesis initiates but is not completed. Using a combination of recombinant and deletion mapping and whole genome sequencing, we identified F09E8.1 as *spe-43*. SPE-43 is predicted to exist in two isoforms; one isoform appears to be a single-pass transmembrane protein while the other is predicted to be a secreted protein. SPE-43 can bind to other known sperm proteins, including SPE-4 and SPE-29, which are known to impact spermiogenesis. In summary, we have identified a membrane protein that is present in *C. elegans* sperm and is required for sperm activation via the hermaphrodite activation signal.

## INTRODUCTION

To fertilize an egg, most sperm must go through the processes of post-meiotic differentiation and activation whereby they gain polarity, motility, and fertilization competence. In *C. elegans* these processes occur concurrently and are referred to collectively as sperm activation or spermiogenesis. During spermiogenesis, round immotile spermatids transition to functional spermatozoa through the formation of a pseudopod, fusion of membranous organelles (MOs) with the plasma membrane, and sorting of proteins and organelles to either the pseudopod or the cell body (Ward et al., 1981; Ward et al., 1983). The initiation of spermiogenesis relies on extracellular signals, allowing for the timing to be precisely controlled in a sex-specific manner (Ellis, 2017). In addition, this signal is relayed through the spermatid without the synthesis of new RNAs or proteins (Ward, 1986). Thus, the study of *C. elegans* sperm activation examines many basic aspects of cell biology including cell signaling, changes in cellular physiology, and establishment of cell polarity through post-translational mechanisms.

Two separate pathways have been identified for the activation of *C. elegans* sperm *in vivo* (Ellis and Stanfield, 2014). Activation of hermaphrodite self-sperm is dependent on the “*spe-8* pathway”. To date, this pathway includes five genes: *spe-8*, *spe-12*, *spe-19*, *spe-27*, and *spe-29*. The *spe-8* gene encodes a non-receptor tyrosine kinase, but the other four genes lack any identifiable functional domains. Mutations in any of these genes result in the same general phenotype: hermaphrodites do not produce self-progeny but males are fertile (Geldziler et al., 2005; Minniti et al., 1996; Muhlrad et al., 2014; Nance et al., 2000; Nance et al., 1999). The extracellular trigger that signals through the *spe-8* group is not yet known. However, a serine protease, TRY-5, present in male seminal fluid was shown to activate sperm through a *spe-8*-independent pathway (Smith and Stanfield, 2011). Thus, male fertility is retained in the absence of any of the *spe-8* group components.

The current model is that SPE-8, SPE-12, SPE-19, SPE-27, and SPE-29 work together in a complex at the sperm plasma membrane where they receive and transduce an extracellular sperm activation signal (Ellis and Stanfield, 2014). This model is supported by data showing that SPE-8 is localized to the spermatid cell membrane and this localization is dependent on *spe-12*, *spe-19*, and *spe-27* (Muhlrad et al., 2014). Additionally, SPE-12, SPE-19, and SPE-29 all have transmembrane domains and SPE-12 has been shown to exist on the sperm cell surface (Nance et al., 1999).

Here, we describe a new gene that is required for sperm activation. We find that *spe-43* displays a classic *spe-8* group phenotype where hermaphrodites are self-sterile due to defective spermiogenesis and males are fertile. Like *spe-12, spe-19*, and *spe-29*, the *spe-43* gene encodes a single-pass transmembrane protein and likely acts as a signaling component within the sperm to initiate sperm activation (Geldziler et al., 2005; Nance et al., 2000; Nance et al., 1999). The SPE-43 protein also contains a DX domain of currently unknown function. Binding studies show that SPE-43 can interact with the *spe-8* group member SPE-29, as well as other sperm proteins with roles in spermiogenesis, sperm activation, and sperm function. Our discovery of *spe-43* adds a new factor to an emerging pathway that cannot be fully understood without the full inventory of components.

## MATERIALS AND METHODS

### Strains and culture methods

General maintenance and crosses of *C. elegans* were performed as described previously (Brenner, 1974). Nematodes were reared at 20°C unless otherwise indicated. Full descriptions of all genotypes used can be found at wormbase.org. Wild type refers to Bristol N2. The Hawaiian strain CB4856 was used for SNP mapping and creation of Hawaiian hybrids for whole-genome sequencing. The *spe-43(eb63)* strain was identified in an ethyl methanesulfonate (EMS) mutagenesis screen for genes with fertility defects by Dr. Andrew Singson while in the L'Hernault laboratory at Emory University. The *spe-43(jn5)* strain was identified in the laboratory of Dr. Gillian Stanfield in an EMS screen for suppressors of *swm-1* (A.M. and G.M.S., unpublished).

### Progeny/ovulation counts

To determine brood sizes and ovulation rates, L4 hermaphrodites were picked onto individual plates and allowed to self-fertilize. Hermaphrodites were transferred to fresh plates daily until new eggs and/or oocytes were no longer observed. Oocytes and/or progeny were counted daily. To assess male fertility, young adult males were crossed to L4 *dpy-5(e61)* hermaphrodites at 20°C in a 4:1 ratio. After 48 hours, the F_0_ worms were removed from the plates. The number of outcross (non-Dpy) and self (Dpy) progeny were counted once the progeny were old enough to score for the Dpy phenotype. Hermaphrodites that died or went missing from the plate were excluded from the analysis. All errors reported are given as standard error of the mean.

### Light microscopy

Differential interference contrast microscopy (DIC) images of live worms or dissected worms were obtained using a Zeiss Universal microscope and captured with an Optronics microscope camera using Magnafire image software (Karl Storz Industrial - America, Inc. El Segundo, CA) or a ProgRes camera (Jenoptik) using ProgResCapturePro software.

### In vitro sperm activation

Males were picked at the L4 or young adult stage and isolated from hermaphrodites for 24 hours. After 24 hours the reproductive tract was dissected in pH 7.8 sperm medium (SM) both with and without the known in vitro activators Pronase (200 μg/ml) or triethanolamine (TEA, 120 mM at pH 7.8) (L'Hernault S and Roberts, 1995; Ward et al., 1983). Hermaphrodites were picked at the L4 stage and isolated from males for 24 hours after which the reproductive tract was dissected in pH 7.8 sperm medium with and without Pronase.

### DAPI staining

To examine sperm cell nuclei, adult hermaphrodites were fixed in cold methanol for 30 seconds then placed in VECTASHIELD antifade mounting medium with DAPI (DAPI concentration is 1.5 ug/ml)(Vector Laboratories, Burlingame, CA) and mounted on 2% agarose slides for viewing using both fluorescence and Nomarski imaging.

### Transactivation assay

*spe-43* L4 hermaphrodites were crossed to *fer-1(hc1);him-5(e1490)*males in a 1:5 ratio at 24.5°C, a temperature at which *fer-1(hc1)* worms are sterile due to a sperm motility defect (Shakes and Ward, 1989; Ward and Miwa, 1978). As controls, *fer-1(hc1); him-5(e1490)* males were crossed to *fer-1(hc1); him-5(e1490)* hermaphrodites. After 48 hours the parents were removed from the plates. The number of progeny produced by each hermaphrodite during that 48 hour period was counted after an additional 2 days.

### spe-6 suppression

Epistasis experiments were performed using the weak hypomorphic allele *spe-6*(*hc163*) as described by (Geldziler et al., 2005; Muhlrad and Ward, 2002). Homozygous *spe-6(hc163) dpy-18(e364)* hermaphrodites were mated to either *spe-43(eb63)* males or *fog-2* males. *fog-2* was used as a control hermaphrodite-specific sterile mutation that cannot be suppressed by *spe-6(hc163).* F1 heterozygotes were allowed to self-fertilize. F2 Dpy hermaphrodites were then picked and scored for sterility. To ensure that the homozygous *spe-43* class of animals was present, individual F_2_ Dpy hermaphrodites were then crossed to homozygous *spe-43* males and the progeny were scored for sterility. Animals for whom all progeny were sterile (2 of 19) indicated that the *spe-43* homozygous class of animal was present.

### Genetic mapping

Linkage, three-factor, and deficiency mapping were carried out as described in (Fay, 2013). Linkage analysis placed *spe-43* on chromosome IV. Three factor mapping further localized *spe-43* to the right arm of chromosome IV. Briefly, *spe-43(eb63)* males were crossed to the strain MT4150 *(unc-17; dpy-4)*, recombinants from among F1 progeny were isolated, and the percent of recombinants was determined. Analysis of the recombinants placed *spe-43* between *unc-17* and *dpy-4*. Using deficiency mapping, we further narrowed down the *spe-43* genetic region. Male *spe-43(eb63)* worms were mated in a 4:1 ratio with hermaphrodites from two different deficiency lines: *sDf21and sDf22.* Heterozygous broods from the different crosses were isolated and scored for the *spe-43* infertility phenotype. *sDf21*and *sDf22* failed to complement *spe-43*, consistent with other mapping data.

### Whole genome sequencing

*spe-43* hermaphrodites were crossed with males from the polymorphic wild-type Hawaiian strain CB4856. F_1_ heterozygous hermaphrodites were crossed with sibling F_1_ heterozygous males in a ratio of 3:15. Individual F_2_ hermaphrodites were picked at the L4 stage to 15 24-well plates and scored the following day for sterility. Sterile (*spe-43* homozygous) hermaphrodites were then crossed in a 1:1 ratio with sibling F_2_ males (of unknown genotype). F_3_ progeny were checked for hermaphrodite self-sterility to identify homozygous *spe-43* male/hermaphrodite lines. Thirteen individual homozygous lines were recovered. F_3_s and F_4_s from these lines were pooled for preparation of genomic DNA by standard phenol/chloroform extraction.

<u>Library construction:</u> Genomic DNA (3 µg of genomic DNA in 110 µl 10 mMTris-Cl, pH 8.5) was sheared to 200-400-bp fragments by using a Bioruptor and 1.5ml Bioruptor Plus TPX tube (Diagenode). Bioruptor was set at the high intensity level with on=0.5 minute and off=0.5 minute for three 10-minute cycles. DNA was precipitated by adding 15.4 μl of 7.5M ammonium acetate, 2 μl glycoblue, and 250 μl of ethanol, and then was pelleted by centrifugation at 20,000xg for 15 minute at room temperature. DNA was dissolved in 20 μl nuclease-free water. The level of fragmentation was confirmed by agarose gel analysis. 10 ng of fragmented DNA was used to prepare the DNA library by using the KAPA Hyper Prep Kit (KAPA Biosystems) as instructed by the manufacturer. 300-500 bp library DNA was selected by using the QIAQuick Gel Extraction Kit (Qiagen) and sequenced on an Illumina HiSeq2000 instrument using the 50-cycle, single-end mode. 82,019,574 reads were obtained.

<u>Sequence alignment:</u> We first collapsed the reads with identical sequences. The resulting 63.2 million unique sequences were aligned to the *C. elegans*N2 strain genome (version WS220) using the software Bowtie (version 0.12.7) (Langmead et al., 2009). 58.6 million reads had at least one reported alignment, either with a perfect match or with a single-nucleotide mismatch. This gives an approximately 27.5x genomic coverage. Only the unambiguously mapped reads were used for SNP analysis.

<u>SNP analysis:</u> 103,346 annotated SNPs between the N2 strain and Hawaiian (HA) strain were used for analysis (downloaded from the CloudMap service on the Galaxy web platform) (Minevich et al., 2012). The average SNP density is approximately 1.0 per kilobase. Among the 2,630,860 reads that covered the SNP locations (on average 25.5 reads per SNP), 1,687,633reads corresponded to the N2 strain and 943,227 toHA (N2:HA = 1.8:1). SNPs covered with more than 12 reads (N2 and HA combined) were used for subsequent analysis.The Hawaiian allele frequency for each SNP, calculated as HA reads/(HA reads+N2 reads), was plotted as a function of genomic location for each chromosome (Figure 4A).

<u>Mutation identification:</u> 72 putative homozygous single-nucleotide substitutions on chromosome IV were identified based on the WGS data using a custom python script (S.G. Gu, unpublished). Each case was supported by at least six different reads with the same single-nucleotide substitution and the lack of wild-type N2 reads.

### Preparation of transgenic rescue lines

Young adult N2 hermaphrodites were microinjected with fosmids WRM061cC06 or WRM0629cE02 (10 µg/ml) along with *myo-3::gfp* (100 μg/ml) as a transformation marker. Worms were also transformed with a PCR product of F09E8.1 (plus 6,014 bp upstream and 169 bp downstream) and *myo-2::gfp* as a transformation marker. The PCR product was amplified from WRM061cC06 using primers 5' - GGA CCG GAA ATT TGA ATA GAA TAG TG and 5' - CCT AAG CCC GGT TTC AGA AT. Once stable transgenic lines were established, they were crossed into the *spe-43(eb63)* mutant background. To show that the transgenic lines were homozygous for *spe-43(eb63)* non-glowing hermaphrodites were plated and confirmed to be sterile. To check for rescue, GFP-positive *spe-43(eb63)* hermaphrodites were scored for self-fertility.

### RT-PCR

RNA was extracted from *him-5(e1490)* and *spe-43(jn5)* worms using TRI Reagent (Molecular Research Center, Inc.) and standard phenol/chloroform extraction protocols. Following extraction, the RNA was treated with DNase to remove any contaminating gDNA. cDNA was then synthesized using an iScript Select cDNA synthesis kit (Bio-Rad).

### Yeast 2-hybrid analysis

The split-ubiquitin yeast two-hybrid assay (Lentze and Auerbach, 2008) was carried out using the DUALmembrane kit (Dualsystems Biotech, USA) as per the manufacturer's instructions. The b isoform (WormBase.org) of the *spe-43* gene was amplified from a male cDNA library and cloned into the bait plasmid pBT3-STE. The details describing the constructs for the prey proteins will be published elsewhere (Marcello *et al.*, in press).

The yeast strain NMY51 was first transformed with SPE-43 bait plasmid and selected on synthetic medium lacking leucine. The strains transformed with bait plasmid were transformed with prey plasmid and were selected on synthetic medium lacking leucine and tryptophan. The resultant strains harboring both bait and prey plasmids were examined for their ability or inability to turn on the reporter genes by plating them on synthetic medium lacking Trp, Leu, His and Ade. Yeast cells transformed with bait plasmid and pAI-Alg5, which expresses wild-type Nub (N-terminal ubiquitin), served as a positive control. Yeast cells transformed with bait plasmid and empty prey vectors pPR3-STE or pPR3-N were used as negative controls.

## RESULTS

### *spe-43* hermaphrodites are sterile

Initial observation of *spe-43(eb63)* showed that hermaphrodites have defects in fertility. The brood size for an individual *spe-43(eb63)* hermaphrodite allowed to self-fertilize is essentially zero at 16°C, 20°C, and 25°C (Figure 1A and data not shown). To determine the ovulation rate of these hermaphrodites, we counted the number of unfertilized eggs laid over the lifespan of the worm. Despite the lack of progeny, the overall ovulation rate for *spe-43(eb63)* is similar to that of wild type (Figure 1B). This result indicates that *spe-43(eb63)* does not interfere in the production of oocytes or the development and function of the somatic gonad. Wild-type rates of ovulation also suggest the presence of spermatids or spermatozoa, which are necessary to stimulate high levels of both oocyte maturation and ovulation (McCarter et al., 1999; Miller et al., 2001).

**Figure 1.**
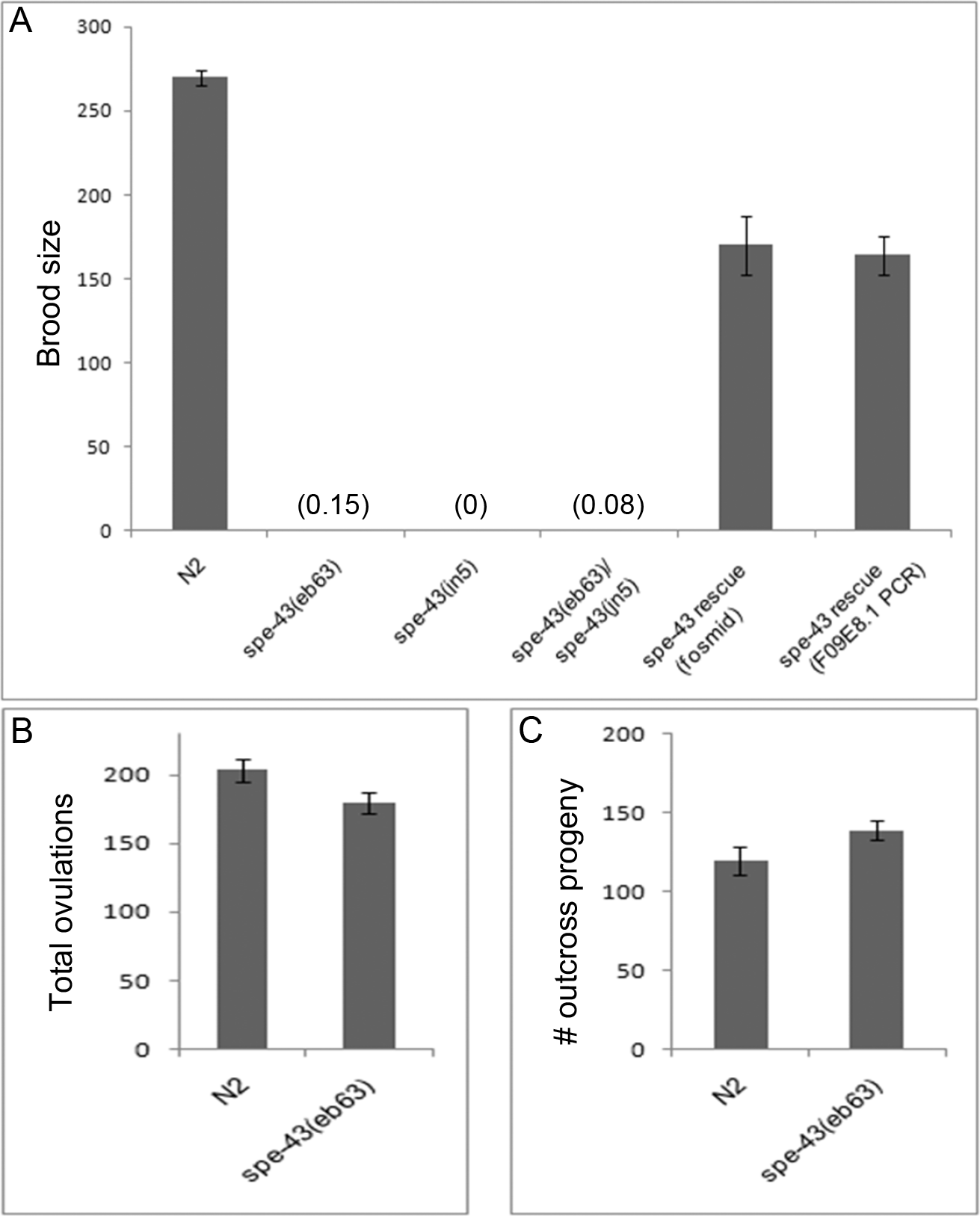
*spe-43* fertility. (A) Hermaphrodite self-fertility at 20°C. Bars show the average number of total self-progeny produced by hermaphrodites. Extra-chromosomal arrays with the fosmid WRM0629cE02 or F09E8.1 PCR product restore self-fertility to *spe-24(eb63)* hermaphrodites, n ≥ 10 (B) Total ovulations is determined by combining the number of self-progeny with the number of unfertilized oocytes laid, n > 15 (C) Male fertility. Males were crossed with *dpy-5* hermaphrodites for 48 hours. Non-Dpy progeny were scored as outcross progeny, n = 20. All error bars represent SEM.

Direct observation of the hermaphrodite adult gonad indicated that *spe-43(eb63)* animals have normal gonad morphology and that pre-fertilization events such as the increase in oocyte size and disappearance of the nucleolus proceed as in wild type. However, their uteri were filled with unfertilized eggs instead of the developing embryos that are present in wild-type animals (Figure 2A, 2B). DAPI staining also confirmed the presence of sperm, their compact nuclei seen as bright puncta in the spermathecae of young adult hermaphrodites (Figure 2C, 2D). However, the sperm were not restricted to the spermathecae and were often seen in the uterus as well (Figure 2D). Since wild-type sperm rapidly crawl back to the spermatheca if they are displaced by a passing oocyte (Ward and Carrel, 1979), the frequent presence of sperm in the uteri of *spe-43(eb63)* animals suggests a defect in sperm motility or migration. When we looked at the reproductive tract of *spe-43(eb63)* hermaphrodites at day 2 of adulthood we observed few, if any, sperm in the spermathecae or uterus, consistent with rapid sperm loss due to a motility defect (Figure 2E, 2F).

**Figure 2.**
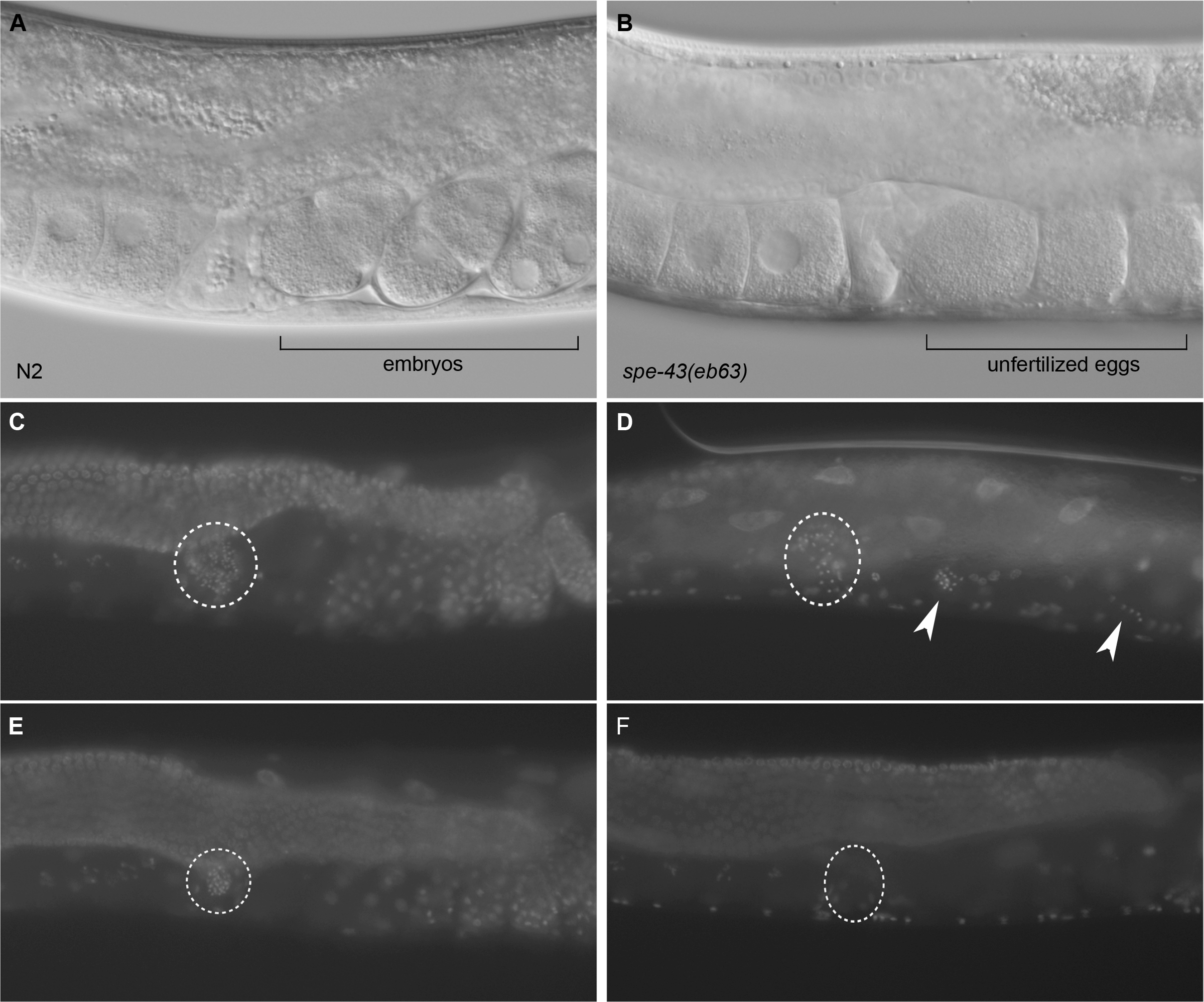
DIC images of N2 (A) and *spe-43(eb63)* (B) hermaphrodite reproductive tracts. N2 hermaphrodites possess developing embryos in the uterus while *spe-43* hermaphrodites have only unfertilized eggs. (C-F) DAPI sperm loss phenotype. DAPI images of 1-day adult N2 (C) and *spe-43(eb63)* (D) hermaphrodites and 2-day adult N2 (E) and *spe-43(eb63)* hermaphrodites grown at 20°C. The position of the spermatheca is marked by dashed circles. Sperm nuclei are visible as small, bright puncta. In *spe-43* mutants, spermatids unable to migrate back into the spermatheca can be found in the uterus and near the vulva (arrow heads).

### *spe-43* males are fertile

Given that the *spe-43(eb63)* hermaphrodite sterility appeared to be caused by a sperm defect, we next sought to examine the impact of *spe-43(eb63)* on male fertility. To measure the fertility of *spe-43(eb63)* males, we crossed *spe-43(eb63)* males to *dpy-5* hermaphrodites and counted the number of outcross (non-Dpy) progeny. The number of outcross progeny produced in a 48 hour period was not significantly different between N2 and *spe-43(eb63)* males (Figure 1C); nor was the total number of outcross progeny produced given a 48 hour mating period (N2 = 245, *spe-43(eb63)* = 252). From these data we can conclude that sperm from *spe-43(eb63)* males activate normally, correctly migrate to the hermaphrodite spermathecae, efficiently out-compete hermaphrodite self-sperm for access to the egg, and have normal fertilization competence. We were also able to establish a *spe-43(eb63)* homozygous line that could be maintained by the presence of *spe-43(eb63)* males. Knowing that *spe-43(eb63)* hermaphrodites do not produce self-progeny, the fertile *spe-43(eb63)* male/hermaphrodite line confirms that a) *spe-43(eb63)* males are fertile and b) *spe-43(eb63)* hermaphrodites can produce viable progeny when supplied with sperm from a fertile male.

### *jn5* is a second allele of spe-43

The *jn5* mutation was isolated in an independent screen for suppressors of *swm-1* (Stanfield and Villeneuve, 2006). This is consistent with previous work that found *spe-8* group activity is required for the ectopic spermatid activation seen in *swm-1* males (Stanfield and Villeneuve, 2006). After outcrossing the *jn5* allele three times to wild type N2 to separate it from the *swm-1* and *him-5* mutations we found that *jn5* males were fertile but *jn5* hermaphrodites were self-sterile (Figure 1A and data not shown). When we crossed *jn5* with *eb63*, the F_1_ trans-heterozygous hermaphrodites remained self-sterile (Figure 1A). The failure of *jn5* to complement *eb63* leads us to conclude that *jn5* is a second allele of *spe-43.*

### *spe-43* spermatids show defects in spermiogenesis

The hermaphrodite-specific sterility and the ability to suppress the sterility of*swm-1* males both point to *spe-43* playing a role in sperm activation through the *spe-8* pathway (Ellis and Stanfield, 2014). To examine the ability of *spe-43* sperm to activate, we dissected sperm from males into sperm media with and without Pronase. In the absence of Pronase, *spe-43* spermatids looked normal and were morphologically indistinguishable from wild type (Figure 3A-C). In the presence of Pronase - a potent *in vitro* activator of *C. elegans* sperm (Ward et al., 1983) - wild-type spermatids rapidly transform into active spermatozoa with dynamic pseudopods (Figure 3D). In contrast, *spe-43* sperm arrested with a “spiked intermediate” phenotype characteristic of mutations in the *spe-8* class of genes (Figure 3E, 3F). If we instead activated the sperm with the weak base TEA, *spe-43* sperm fully activated and extended pseudopods in a manner comparable to wild type (data not shown). This ability to activate in TEA, despite the failure to activate with Pronase, is true for most members of the *spe-8* group (Minniti et al., 1996; Nance et al., 2000; Shakes and Ward, 1989).

**Figure 3.**
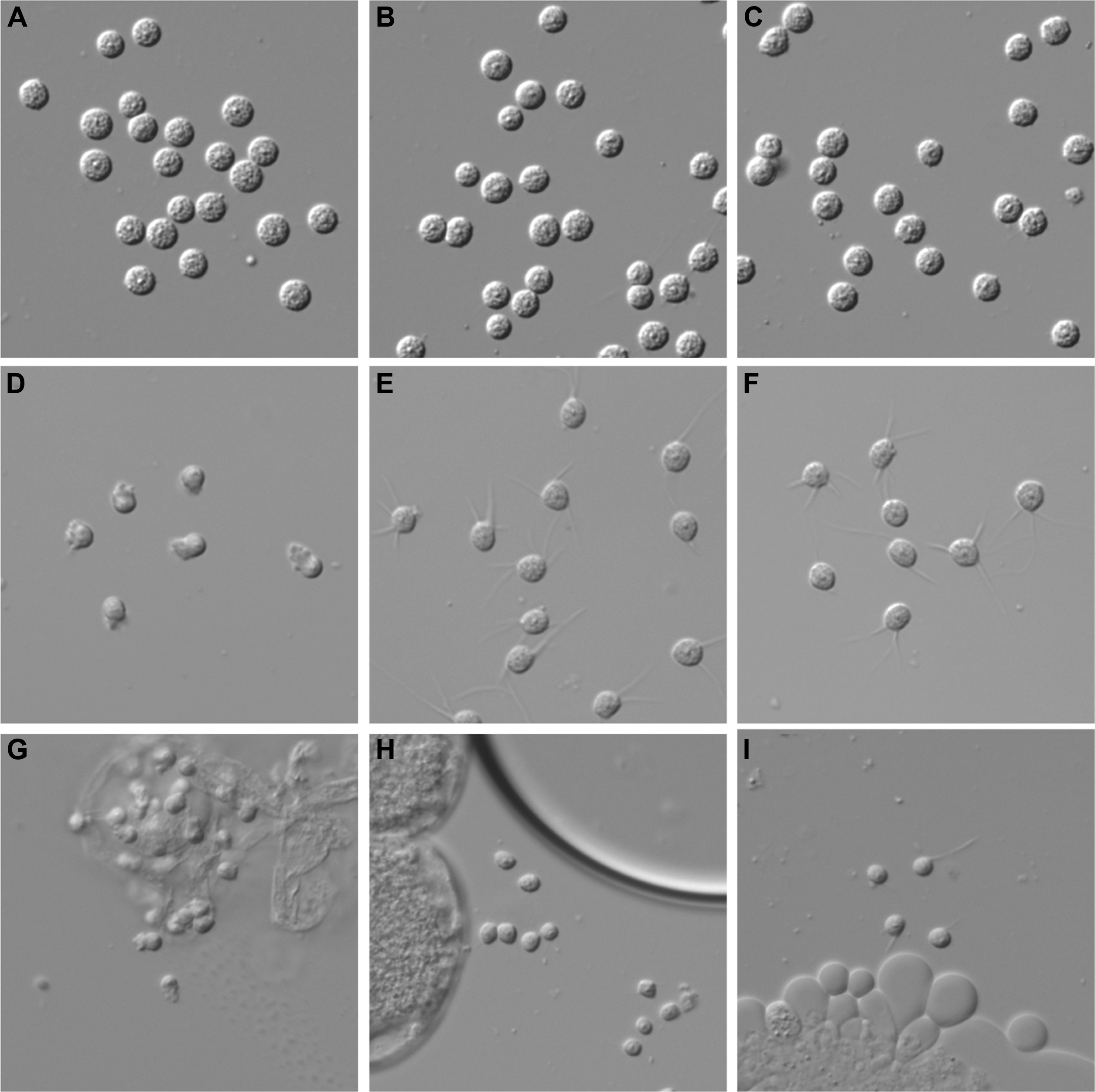
Sperm activation phenotypes. Spermatids from adult males were dissected in sperm medium (A-C) or sperm medium with Pronase (D-F). (A, D) N2; (B, E) *spe-43(eb63);* (C, F) *spe-43(jn5)*. (G-I) Spermatids from young adult hermaphrodites. (G) N2 hermaphrodite sperm in sperm medium. (H) *spe-43(eb63)* hermaphrodite sperm in sperm medium. (I) *spe-43(eb63)* hermaphrodite sperm in sperm medium with Pronase.

We also looked at hermaphrodite sperm by dissecting unmated one day adult N2 and *spe-43* hermaphrodites. Normally, hermaphrodite sperm activate when they enter the spermathecae (Ward and Carrel, 1979) and no *in vitro* activator is required for pseudopods to be observed (Figure 3G). We were unable to find any activated sperm when we dissected *spe-43* hermaphrodites, and instead found only spermatids (Figure 3H). Like *spe-43* male spermatids, *spe-43* hermaphrodite spermatids arrested at the spiked intermediate stage of activation if Pronase is added to the sperm media (Figure 3I).

Finally, mutations that exhibit activation defects only in the hermaphrodite are sometimes capable of the phenomenon known as "transactivation", whereby TRY-5 transferred in the seminal fluid of males can activate the hermaphrodite self-sperm (Minniti et al., 1996; Nance et al., 2000; Nance et al., 1999; Shakes and Ward, 1989; Smith and Stanfield, 2011). We tested whether *spe-43* hermaphrodite sperm could be transactivated by crossing *fer-1(hc1); him-5(e1490)* males to *spe-43(eb63)* and *spe-43(jn5)* hermaphrodites. The *fer-1* mutation leads to defective sperm that cannot fertilize, but these males still succeed in transferring seminal fluid to the hermaphrodite (Shakes and Ward, 1989; Ward and Miwa, 1978). Therefore, any progeny produced from this cross are self-progeny of the hermaphrodite resulting from activation of the hermaphrodite's self-sperm through the male activation pathway. We observed a small amount of transactivation for both *spe-43* alleles (*eb63* = 4.1 progeny, n=15; *jn5* = 9.9 progeny, n=15; fer-1 = 0 progeny, n=15). These levels are similar to previously reported levels of transactivation for other severe loss-of-function activation mutants (Geldziler et al., 2005; Nance et al., 1999). Although the number of progeny from transactivation was low, these results suggest that *spe-43* hermaphrodite sperm are competent to undergo spermiogenesis. Thus we conclude that SPE-43 is required for response to the hermaphrodite activator and not for the cellular changes of sperm activation per se.

### *spe-6(hc163)* suppresses spe-43 sterility

To confirm that *spe-43* acts in the *spe-8* pathway, we tested the ability of the hypomorphic allele *spe-6(hc163)* to suppress *spe-43* hermaphrodite self-sterility. The *hc63* allele of *spe-6* has previously been shown to suppress the sterility associated with all other *spe-8* class genes: *spe-8, spe-12, spe-19, spe-27*, and *spe-29* (Geldziler et al., 2005; Muhlrad and Ward, 2002). We crossed *spe-6(hc163) dpy-18(e364)* hermaphrodites to *spe-43(eb63)* males and scored F_2_ Dpy hermaphrodites for self-fertility. Approximately one-fourth of the F_2_ Dpys should be sterile unless *spe-6* is able to suppress *spe-43(eb63)*. We observed that all 210 Dpy hermaphrodites tested were fertile, indicating that *spe-6(hc163)* does indeed suppress *spe-43(eb63)*. As a control, we also crossed *spe-6(hc163) dpy-18(e364)* hermaphrodites to *fog-2(q71)* males, since *fog-2* is not a component of the *spe-8* pathway, and observed the expected sterile F_2_ Dpys (42/143). Thus, we conclude that *spe-43* functions upstream of *spe-6* during *C. elegans* spermiogenesis and is a component of the same genetic pathway as *spe-8* and the other *spe-8* class genes.

### Identification of the *spe-43* gene

We took a mapping-by-sequencing approach to determine the causative mutation for the *spe-43(eb63)* phenotype (Doitsidou et al., 2010) (Materials and Methods). This strategy identified an approximately 2.4 Mbp region on chromosome IV (13,100,000 - 15,500,000 bp) that showed a strong reduction in Hawaiian SNP frequency (Figure 4A). This region matched the predicted location of *spe-43(eb63)* based on traditional mapping data (see Materials and Methods) making it the target region for further analysis of the whole genome sequencing data. Among the putative homozygous single-nucleotide substitutions within this region, the only nonsense mutation (chrIV:13,141,921 C → T, amber mutation) was mapped to the 3rd exon of the protein-coding gene *F09E8.1*. Sanger sequencing confirmed the presence of this mutation in the *spe-43(eb63)* strain but not in the lab N2 strain. Because of its genomic location and sperm-enriched mRNA expression (Reinke et al., 2000), *F09E8.1* was further investigated as a candidate for *spe-43*.

**Figure 4.**
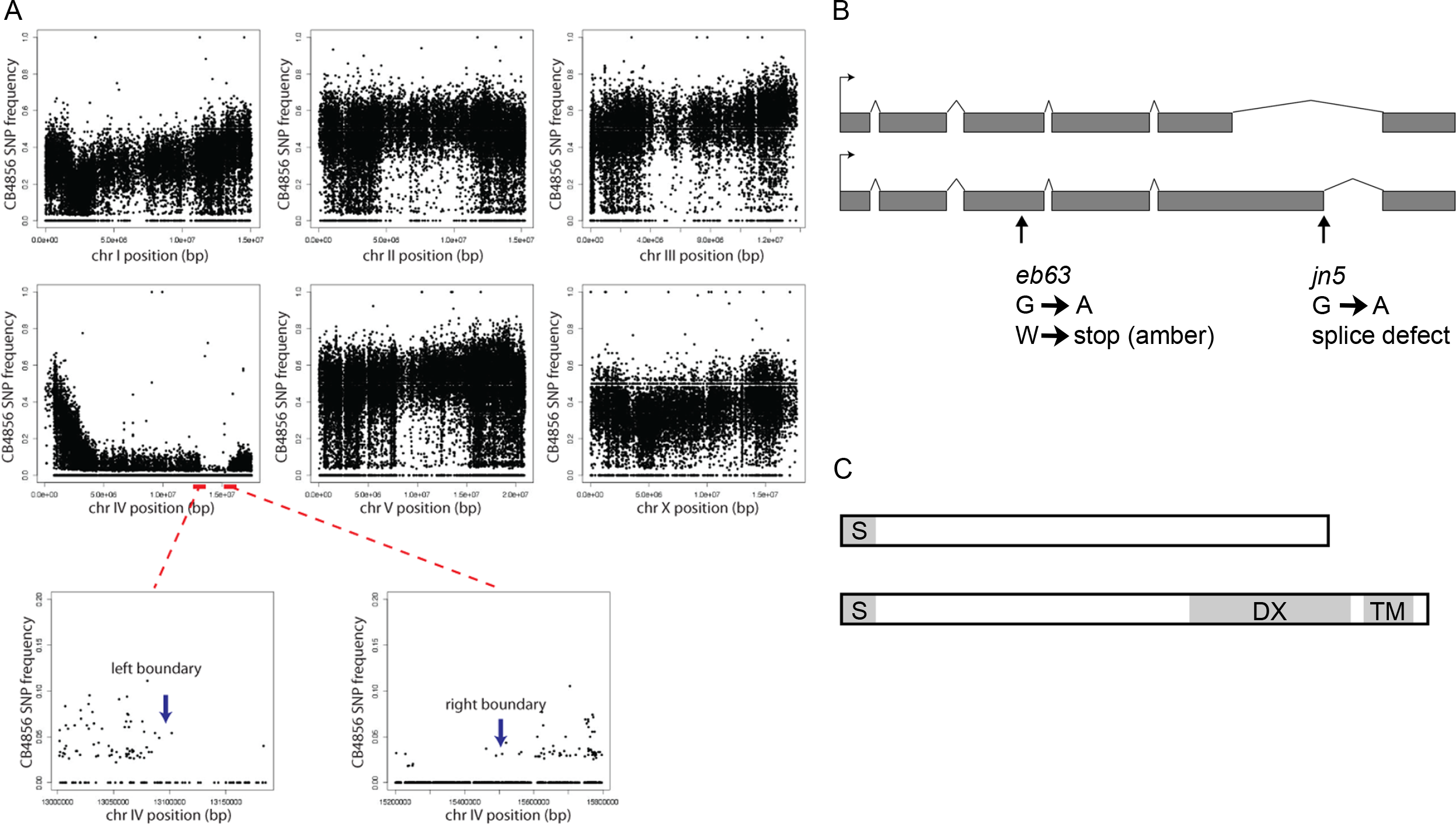
Molecular identity of *spe-43*. (A) Mapping data using Hawaiian hybrids and whole genome sequencing. *spe-43(eb63)* is in the region of chromosome IV where Hawaiian (CB4856) SNP frequency approaches zero. (B) Predicted *spe-43* gene structure (Wormbase). The gene is transcribed as an a isoform (top) and a b isoform (bottom). The location of the mutation associated with each allele is indicated. (C) Predicted protein structures of the a isoform (top) and b isoform (bottom) of SPE-43. Alternative splicing of the fifth exon of the transcript leads to a frameshift of the sixth exon so that the two protein isoforms have unique amino acid sequences after the splice site. S = signal peptide; DX = domain of unknown function DX; TM = transmembrane domain.

We created transgenic animals carrying extrachromosomal arrays containing the fosmid WRM061cC06 or WRM0629cE02, both of which include the full *F09E8.1* genomic sequence, and tested for the ability to rescue *spe-43* sterility. We measured *spe-43(eb63)* self-broods for two independent lines carrying WRM0629cE02 and one line carrying WRM061cC06 and found that both fosmids were able to restore hermaphrodite fertility (Figure 1A). Additionally, we found that a PCR product of *F09E8.1* (plus upstream and downstream sequence to the boundaries of neighboring genes) was able to rescue *spe-43(eb63)* fertility at levels equivalent to what we observed with the fosmids (Figure 1A). From this, we conclude that *spe-43* is *F09E8.1*.

The *spe-43* transcript is composed of six exons and predicted to exist in two isoforms produced by alternative splicing of the fifth exon (Figure 4B). Using cDNA from a mixed population of *him-5* hermaphrodites and males we observed two bands by RT-PCR that corresponded to the predicted isoforms, indicating that both transcripts are made (data not shown). The a and b isoforms are predicted to encode 226 and 273 amino acid proteins, respectively (Figure 4C). Analysis of the protein sequences suggests that both contain a signal sequence at the N-terminus (SignalP). While the a isoform has no other identifiable domains, the b isoform additionally contains a "DX" domain as well as a predicted transmembrane domain that places only the final six C-terminal amino acids in the cytoplasm of the sperm [SMART (Letunic et al., 2015); TMHMM v2.0].

When we sequenced the *spe-43(jn5)* line we found only a silent mutation in the coding region of *F09E8.1*. Yet, the non-complementation between *eb63* and *jn5* led us to investigate how this silent mutation could cause the *spe-43(jn5)* phenotype. We discovered that while the mutation does not alter the predicted protein sequence, it changes the last base of the fifth exon of the b isoform from a G to an A and we predicted that this might alter splicing of the transcript (Figure 4B). RT-PCR of *F09E8.1* confirmed that the b isoform of the transcript is absent in *spe-43(jn5)* worms (Supplementary Figure 1). This observation also allows us to conclude that the b isoform of *spe-43* is necessary for sperm activation; although it does not allow us to draw any conclusions about the necessity of the a isoform.

### SPE-43 physically interacts with other sperm proteins

Since the domain architecture of the predicted SPE-43 protein offers little insight into its potential functions, we tested for direct physical interactions with known *C. elegans* sperm proteins to identify components with which SPE-43 might function. Using a membrane yeast-2-hybrid system (Lentze and Auerbach, 2008), we tested the b isoform of SPE-43 as bait against a panel of 19 sperm proteins (including itself). In addition to interacting with itself, SPE-43 positively interacted with six of the other proteins tested (Table 1). These include proteins with known roles in spermatogenesis (SPE-4, SPE-10, and SPE-15), spermiogenesis (SPE-4, SPE-29), and sperm function (SPE-38, SPE-42).

**Table 1.** Interactions between SPE-43 and other sperm proteins.

| Spermatogenesis |  | Spermiogenesis |  | Sperm Function |  |
| --- | --- | --- | --- | --- | --- |
| SPE-4 | ++ | SPE-29 | + | SPE-38 | + |
| SPE-10 | ++ | SPE-43 | + | SPE-42 | + |
| SPE-15 | +/- | SPE-8 | - | SPE-9 | - |
| SPE-5 | - | SPE-12 | - | SPE-41 | - |
| SPE-6 | - | SPE-19 | - | SPE-45 | - |
| SPE-17 | - | SPE-27 | - |  |  |
| WEE-1.3 | - | FER-1 | - |  |  |

Table 1. Interactions between SPE-43 and other sperm proteins. ++ = strong interaction, + = medium interaction, +/− = weak interaction, − = no interaction observed. Strength of interactions was determined by amount of yeast growth on selective medium relative to negative controls.

## DISCUSSION

We have characterized and genetically identified the *spe-43* gene, showing that it functions in *C. elegans* sperm activation as a component of the *spe-8* pathway. Like other members of this pathway, *spe-43* is essential for hermaphrodite self-fertility, but dispensable for male fertility. Failure of *spe-43* spermatids to activate *in vitro* in the presence of Pronase supports a role for SPE-43 within sperm as a transducer or effector of the extracellular hermaphrodite activation signal. Since the fertility of *spe-43* males and the transactivation of *spe-43* hermaphrodite sperm suggest that SPE-43 is not fundamentally required for the cellular changes of sperm activation nor sperm function, we favor the hypothesis that SPE-43 is part of the signal transduction pathway that relays the hermaphrodite activation signal through the cell.

Similar to most members of the SPE-8 group, the domain architecture of SPE-43 does not offer any obvious predictions regarding how it may function as a signal transducer. While orthologs exist in closely related *Caenorhabditis* species, similar proteins do not appear in other phyla. Our RT-PCR data indicate that two isoforms of *spe-43* are transcribed. We conclude from our analysis of *spe-43(jn5)*, which only affects the b isoform, that the b isoform is necessary for *C. elegans* sperm activation via the hermaphrodite activation pathway. However, we do not currently know if the a isoform has any function. Future work will seek to determine if the b isoform is sufficient for SPE-43's role during sperm activation, thus ruling out any role of the a isoform. Only the b isoform has any identifiable domains based on sequence analysis, lending support to the hypothesis that it is the only functional isoform of SPE-43. The DX domain present in the b isoform is a domain of unknown function that is also restricted to nematodes. *C. elegans* has only seven other DX domain containing proteins, six of which are also predicted transmembrane proteins [Pfam (Finn et al., 2016)]. However, SPE-43 is the first DX domain-containing protein for which a phenotype is being reported. More analysis will be required to determine if this domain is critical to SPE-43 function. In addition, a broader analysis of the entire class of DX proteins could lead to a better understanding of the general function of this domain.

The single transmembrane domain of SPE-43 places only the final six C-terminal amino acids in the cytoplasm of the sperm. Thus, one possibility is that SPE-43 is embedded in the sperm plasma membrane and majority of the SPE-43 protein is localized extracellularly, making it a prime candidate as a receptor for an extracellular sperm activation signal. Epistasis analysis with *spe-6(hc163)* lends genetic evidence that *spe-43* acts in the same molecular pathway as *spe-8, spe-12, spe-19, spe-27*, and *spe-29*, as this specific *spe-6* hypomorph bypasses the need for any of these components, including *spe-43*, for hermaphrodite sperm activation. If SPE-43 is a receptor for the proposed SPE-8 complex at the plasma membrane we might predict physical interactions between SPE-43 and the other components of the complex. However, our yeast 2-hybrid analysis detected a positive interaction with SPE-29 but none of the other known spermiogenesis proteins. Interestingly, *spe-29*is the one member of the *spe-8* group that does not appear to be important for SPE-8 localization (Muhlrad et al., 2014). These findings lead us to propose a model in which SPE-43 and SPE-29 work together within the signal transduction pathway separate from SPE-8, SPE-12, SPE-19, and SPE-27. Of course, we cannot rule out the possibility that interactions between SPE-43 and the "SPE-8 complex" are mediated by an, as of yet, unidentified factor and all these components do form one large receptor/signaling complex. Supporting our model where SPE-43 and SPE-29 represent a separate complex is the indication that SPE-43 may localize to the MOs, rather than the plasma membrane, since positive interactions were found with SPE-4, SPE-10, and SPE-38 and all of these proteins are known to localize to the MOs of spermatids (Arduengo et al., 1998; Chatterjee et al., 2005; Gleason et al., 2006). Further investigations, including direct observation of SPE-43 and SPE-29 localization, will be required to determine exactly where and how SPE-43 is functioning in the cell.

Our binding studies also suggest that SPE-43 forms a dimer or multimer which may relate to its function in the sperm. Additionally, we wonder if our binding studies represent a more functional relationship between spermatogenesis, spermiogenesis, and sperm function proteins. Co-localization to the same organelle does not necessitate direct interactions between proteins. So perhaps SPE-10 and SPE-4 bind SPE-43 because they play a role in getting SPE-43 to the MO during spermatogenesis. Likewise, SPE-43 may bind SPE-38 because it plays a role in regulating SPE-38 localization before and after sperm activation. Thus, the ability to activate is being set up during spermatogenesis and spermiogenesis confers sperm function partially through direct interactions between proteins that we typically think of as acting during separate and distinct periods of sperm development.

In summary, we add *spe-43* to the list of genes that act within the*spe-8* pathway for *C. elegans* sperm activation. Knowing the full complement of molecules that comprise this pathway will be necessary to piece together the mechanism by which the cell signals its change in morphology in the absence of transcription or translation. As more components are added, it also becomes less likely that all these proteins are part of the receptor complex that receives the initiating activation signal. Despite the similarity in general phenotype we need to begin teasing apart the receptor(s) from the molecules that relay the signal through the cell and to the effector proteins carrying out the morphological changes. Additional molecular, biochemical, and genetic examinations will be necessary to discover the precise role that each of these proteins plays in this signal transduction pathway.

## ACKNOWLEDGEMENTS

This work was supported by National Institutes of Health R01HD054681 to AS, R01GM087705 to GMS, and 1K12GM093854 that supported ARK. We thank the CGC (which is funded by NIH Office of Research Infrastructure Programs P40 OD010440) and WormBase. We also thank Matt Marcello for useful discussion regarding SPE-43 binding data.

---

**Supplementary Figure 1.**
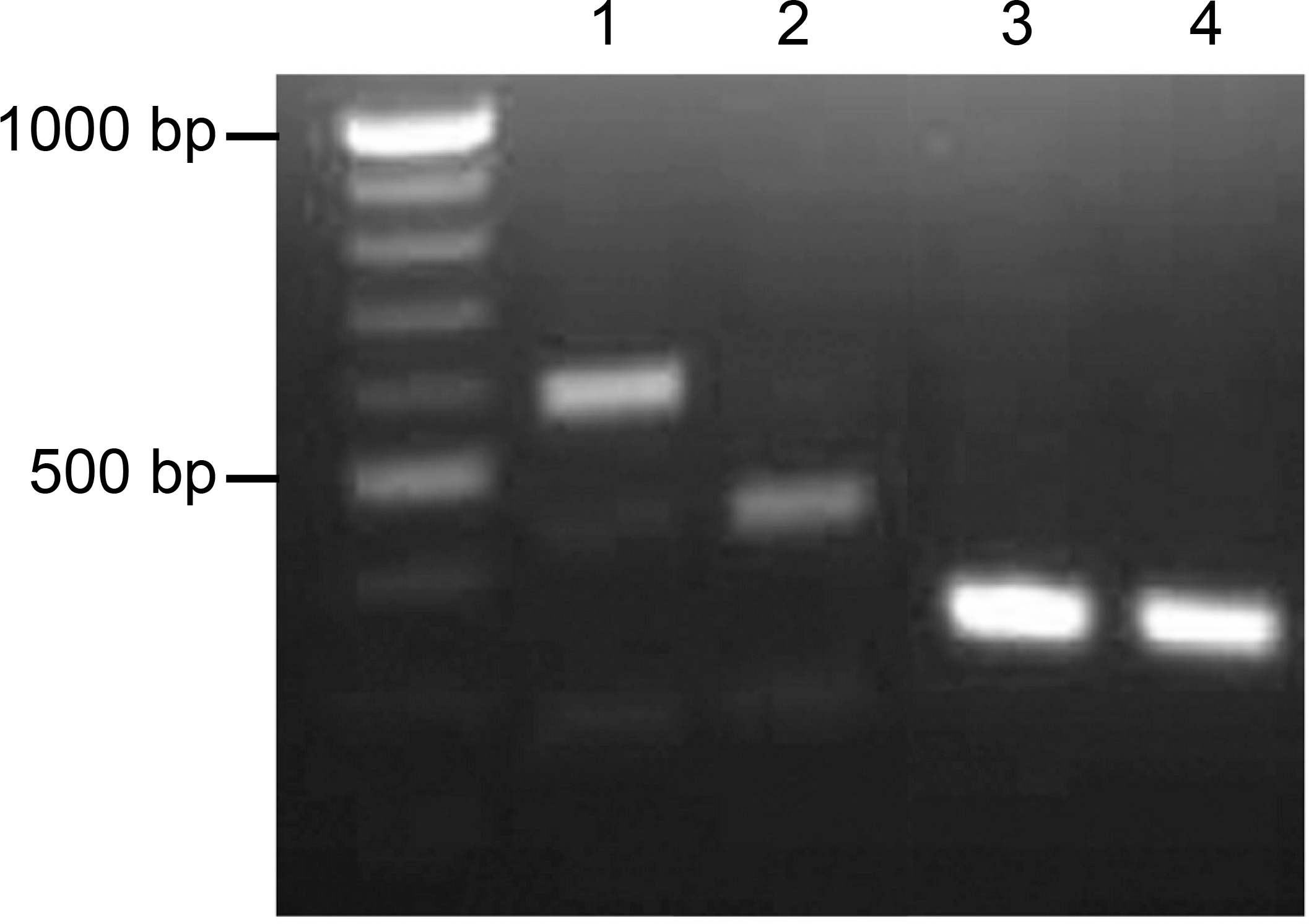
RT-PCR of *spe-43* from *him-5(e1490)* (lane 1) and *spe-43(jn5)* (lane 2). The larger band representing the b isoform is not observed from *spe-43(jn5)* indicating the mutation at the splice site results in a loss of the b isoform transcript in *spe-43(jn5)* animals. The smaller band representing the a isoform is still observed. Control RT-PCR of actin *(act-2)* from *spe-43(jn5)* (lane 3) and *him-5(e1490)* (lane 4).

**Supplementary Figure 2.**
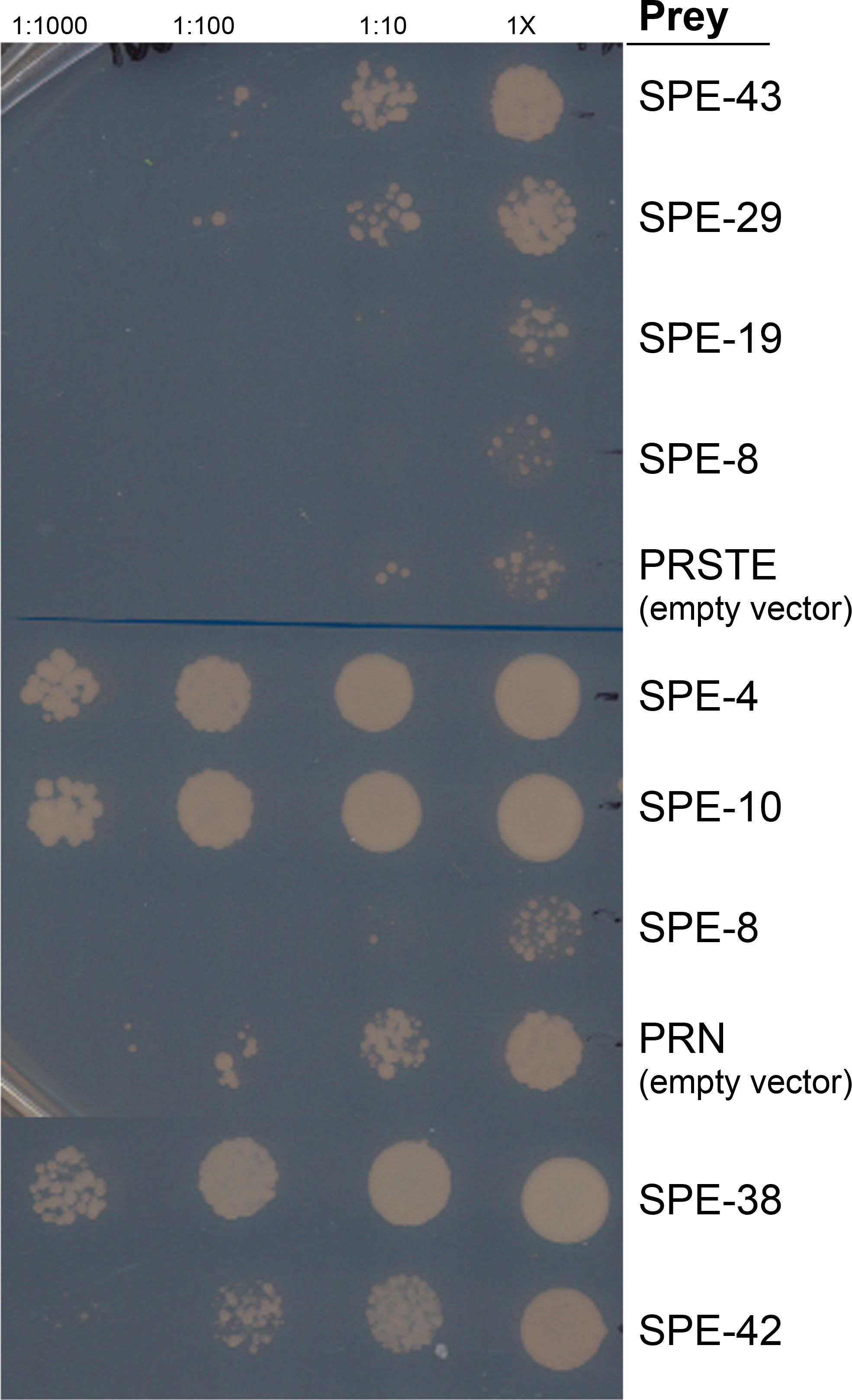
Yeast-2-hybrid interactions with SPE-43 as the bait and various sperm proteins as the prey. Growth indicates a positive interaction between SPE-43 and the prey protein. SPE-43, SPE-29, SPE-19 and SPE-8 (upper) were expressed with the PRSTE vector. SPE-4, SPE-10, SPE-8 (lower), SPE-38, and SPE-42 were expressed with the PRN vector. Empty vectors are negative controls.

## REFERENCES

Arduengo, P. M., Appleberry, O. K., Chuang, P., L'Hernault, S. W., 1998. The presenilin protein family member SPE-4 localizes to an ER/Golgi derived organelle and is required for proper cytoplasmic partitioning during Caenorhabditis elegans spermatogenesis. J Cell Sci. 111 (Pt 24), 3645–54.

Brenner, S., 1974. The genetics of Caenorhabditis elegans. Genetics. 77, 71–94.

Chatterjee, I., Richmond, A., Putiri, E., Shakes, D. C., Singson, A., 2005. The Caenorhabditis elegans spe-38 gene encodes a novel four-pass integral membrane protein required for sperm function at fertilization. Development. 132, 2795–808.

Doitsidou, M., Poole, R. J., Sarin, S., Bigelow, H., Hobert, O., 2010. C. elegans mutant identification with a one-step whole-genome-sequencing and SNP mapping strategy. PLoS One. 5, e15435.

Ellis, R. E., 2017. "The persistence of memory"-Hermaphroditism in nematodes. Mol Reprod Dev. 84, 144–157.

Ellis, R. E., Stanfield, G. M., 2014. The regulation of spermatogenesis and sperm function in nematodes. Semin Cell Dev Biol. 29, 17–30.

Fay, D., Genetic mapping and manipulation: Chapter 1-Introduction and basics. In: T. C. e. R. Community, (Ed.), WormBook. WormBook.

Finn, R. D., Coggill, P., Eberhardt, R. Y., Eddy, S. R., Mistry, J., Mitchell, A. L., Potter, S. C., Punta, M., Qureshi, M., Sangrador-Vegas, A., Salazar, G. A., Tate, J., Bateman, A., 2016. The Pfam protein families database: towards a more sustainable future. Nucleic Acids Res. 44, D279–85.

Geldziler, B., Chatterjee, I., Singson, A., 2005. The genetic and molecular analysis of spe-19, a gene required for sperm activation in Caenorhabditis elegans. Dev Biol. 283, 424–36.

Gleason, E. J., Lindsey, W. C., Kroft, T. L., Singson, A. W., L'Hernault S, W., 2006. spe-10 encodes a DHHC-CRD zinc-finger membrane protein required for endoplasmic reticulum/Golgi membrane morphogenesis during Caenorhabditis elegans spermatogenesis. Genetics. 172, 145–58.

L'Hernault S, W., Roberts, T. M., Cell Biology of Nematode Sperm. In: H. F. Epstein, D. Shakes, Eds.), Caenorhabditis elegans Modern Biological Analysis of an Organism, Vol. 48. Academic Press, Inc., San Diego, 1995, pp. 273–299.

Langmead, B., Trapnell, C., Pop, M., Salzberg, S. L., 2009. Ultrafast and memory-efficient alignment of short DNA sequences to the human genome. Genome Biol. 10, R25.

Lentze, N., Auerbach, D., 2008. Membrane-based yeast two-hybrid system to detect protein interactions. Curr Protoc Protein Sci. Chapter 19, Unit 19 17.

Letunic, I., Doerks, T., Bork, P., 2015. SMART: recent updates, new developments and status in 2015. Nucleic Acids Res. 43, D257–60.

McCarter, J., Bartlett, B., Dang, T., Schedl, T., 1999. On the control of oocyte meiotic maturation and ovulation in Caenorhabditis elegans. Dev Biol. 205, 111–28.

Miller, M. A., Nguyen, V. Q., Lee, M. H., Kosinski, M., Schedl, T., Caprioli, R. M., Greenstein, D., 2001. A sperm cytoskeletal protein that signals oocyte meiotic maturation and ovulation. Science. 291, 2144–7.

Minevich, G., Park, D. S., Blankenberg, D., Poole, R. J., Hobert, O., 2012. CloudMap: a cloud-based pipeline for analysis of mutant genome sequences. Genetics. 192, 1249–69.

Minniti, A. N., Sadler, C., Ward, S., 1996. Genetic and molecular analysis of spe-27, a gene required for spermiogenesis in Caenorhabditis elegans hermaphrodites. Genetics. 143, 213–23.

Muhlrad, P. J., Clark, J. N., Nasri, U., Sullivan, N. G., LaMunyon, C. W., 2014. SPE-8, a protein-tyrosine kinase, localizes to the spermatid cell membrane through interaction with other members of the SPE-8 group spermatid activation signaling pathway in C. elegans. BMC Genet. 15, 83.

Muhlrad, P. J., Ward, S., 2002. Spermiogenesis initiation in Caenorhabditis elegans involves a casein kinase 1 encoded by the spe-6 gene. Genetics. 161, 143–55.

Nance, J., Davis, E. B., Ward, S., 2000. spe-29 encodes a small predicted membrane protein required for the initiation of sperm activation in Caenorhabditis elegans. Genetics. 156, 1623–33.

Nance, J., Minniti, A. N., Sadler, C., Ward, S., 1999. spe-12 encodes a sperm cell surface protein that promotes spermiogenesis in Caenorhabditis elegans. Genetics. 152, 209–20.

Reinke, V., Smith, H. E., Nance, J., Wang, J., Van Doren, C., Begley, R., Jones, S. J., Davis, E. B., Scherer, S., Ward, S., Kim, S. K., 2000. A global profile of germline gene expression in C. elegans. Mol Cell. 6, 605–16.

Shakes, D. C., Ward, S., 1989. Initiation of spermiogenesis in C. elegans: a pharmacological and genetic analysis. Dev Biol. 134, 189–200.

Smith, J. R., Stanfield, G. M., 2011. TRY-5 is a sperm-activating protease in Caenorhabditis elegans seminal fluid. PLoS Genet. 7, e1002375.

Stanfield, G. M., Villeneuve, A. M., 2006. Regulation of sperm activation by SWM-1 is required for reproductive success of C. elegans males. Curr Biol. 16, 252–63.

Ward, S., Asymmetric localization of gene products during the development of Caenorhaditis elegans spermatozoa. In: J. G. Gall, (Ed.), Gametogenesis and the Early Embryo. Alan R. Liss, Inc, New York, 1986, pp. 55–75.

Ward, S., Argon, Y., Nelson, G. A., 1981. Sperm morphogenesis in wild-type and fertilization-defective mutants of Caenorhabditis elegans. J Cell Biol. 91, 26–44.

Ward, S., Carrel, J. S., 1979. Fertilization and sperm competition in the nematode Caenorhabditis elegans. Dev Biol. 73, 304–21.

Ward, S., Hogan, E., Nelson, G. A., 1983. The initiation of spermiogenesis in the nematode Caenorhabditis elegans. Dev Biol. 98, 70–9.

Ward, S., Miwa, J., 1978. Characterization of temperature-sensitive, fertilization-defective mutants of the nematode caenorhabditis elegans. Genetics. 88, 285–303.

